# Evaluation of mRNA-1273 against SARS-CoV-2 B.1.351 Infection in Nonhuman Primates

**DOI:** 10.1101/2021.05.21.445189

**Authors:** Kizzmekia S. Corbett, Anne P. Werner, Sarah O’ Connell, Matthew Gagne, Lilin Lai, Juan I. Moliva, Barbara Flynn, Angela Choi, Matthew Koch, Kathryn E. Foulds, Shayne F. Andrew, Dillon R. Flebbe, Evan Lamb, Saule T. Nurmukhambetova, Samantha J. Provost, Kevin W. Bock, Mahnaz Minai, Bianca M. Nagata, Alex Van Ry, Zackery Flinchbaugh, Timothy S. Johnston, Elham Bayat Mokhtari, Prakriti Mudvari, Amy R. Henry, Farida Laboune, Becky Chang, Maciel Porto, Jaclyn Wear, Gabriela S. Alvarado, Seyhan Boyoglu-Barnum, John-Paul M. Todd, Bridget Bart, Anthony Cook, Alan Dodson, Laurent Pessaint, Katelyn Steingrebe, Sayda Elbashir, Hanne Andersen, Kai Wu, Darin K. Edwards, Swagata Kar, Mark G. Lewis, Eli Bortiz, Ian N. Moore, Andrea Carfi, Mehul S. Suthar, Adrian McDermott, Mario Roederer, Martha C. Nason, Nancy J. Sullivan, Daniel C. Douek, Barney S. Graham, Robert A. Seder

## Abstract

**Background:** Vaccine efficacy against the B.1.351 variant following mRNA-1273 vaccination in humans has not been determined. Nonhuman primates (NHP) are a useful model for demonstrating whether mRNA-1273 mediates protection against B.1.351.

**Methods:** Nonhuman primates received 30 or 100 µg of mRNA-1273 as a prime-boost vaccine at 0 and 4 weeks, a single immunization of 30 µg at week 0, or no vaccine. Antibody and T cell responses were assessed in blood, bronchioalveolar lavages (BAL), and nasal washes. Viral replication in BAL and nasal swabs were determined by qRT-PCR for sgRNA, and histopathology and viral antigen quantification were performed on lung tissue post-challenge.

**Results:** Eight weeks post-boost, 100 µg x2 of mRNA-1273 induced reciprocal ID_50_ neutralizing geometric mean titers against live SARS-CoV-2 D614G and B.1.351 of 3300 and 240, respectively, and 430 and 84 for the 30 µg x2 group. There were no detectable neutralizing antibodies against B.1351 after the single immunization of 30 µg. On day 2 following B.1.351 challenge, sgRNA in BAL was undetectable in 6 of 8 NHP that received 100 µg x2 of mRNA-1273, and there was a ∼2-log reduction in sgRNA in NHP that received two doses of 30 µg compared to controls. In nasal swabs, there was a 1-log_10_ reduction observed in the 100 µg x2 group. There was limited inflammation or viral antigen in lungs of vaccinated NHP post-challenge.

**Conclusions:** Immunization with two doses of mRNA-1273 achieves effective immunity that rapidly controls lower and upper airway viral replication against the B.1.351 variant in NHP.

## INTRODUCTION

The severe acute respiratory syndrome coronavirus-2 (SARS-CoV-2) pandemic has led to more than 3.4 million deaths worldwide^1^. Vaccination with two 100 µg doses of mRNA-1273, a lipid nanoparticle (LNP) encapsulated messenger RNA-based vaccine encoding a stabilized full-length SARS-CoV-2 Wuhan-Hu-1 spike (S) glycoprotein, proved 94% efficacious against symptomatic COVID-19 in the United States (US)^2^. mRNA-1273 is authorized by the US Food and Drug Administration for Emergency Use (EUA).

The emergence of SARS-CoV-2 variants of concern (VOC)^3^ that show reduced neutralization by sera from Wu-1 strain convalescent subjects or vaccinees^4-6^ has created uncertainty about the efficacy of current SARS-CoV-2 vaccines against VOC infection. To date, the most concerning variants contain combinations of mutations and deletions in the S receptor-binding domain (RBD) and N-terminal domain (NTD), respectively. Acquisition of amino acid substitutions in the S RBD-- namely K417N, E484K, and N501Y—and in the NTD, such as L18F, D80A, D215G, and Δ242-244, is associated with increased transmissibility and reduction in neutralization sensitivity^7-17^. Variants containing these substitutions originally isolated in the United Kingdom (UK) (B.1.1.7), Republic of South Africa (B.1.351), Brazil (P.1 lineage), New York (B.1.526), and California (B.1.427/B.1.429), have shown varying reduction in neutralization by convalescent and vaccine serum, and are resistant to some monoclonal antibodies^14,18-24^. Among these variants, B.1.351 contains the most concerning set of mutations in the RBD and NTD subdomains^25^.

We and others recently reported that sera from mRNA-1273-immunized human and nonhuman primates (NHP) showed the greatest reduction of neutralization against B.1.351 compared to B.1.1.7, P.1, B.1.427/B.1.429, and B.1.1.7+E484K variants^7-17,26^. In UK- or US-based clinical studies, NVX-CoV2373 (Novavax), AZD1222 (University of Oxford/AstraZeneca), and Ad26.COV2.S (Janssen/Johnson & Johnson) vaccines show between ∼70 and 90% protection against the circulating D614G or B.1.1.7 variants^11,27-29^, and vaccine efficacy against mild symptomatic COVID-19 caused by B.1.351 was up to 60% for Ad26.CoV2^29^ and NVX-CoV2373^30^ and ∼10% for AZD122^31^. A recent report showed BNT162b2, Pfizer’s mRNA vaccine, conferred ∼75% protection against confirmed B.1.351 infection in Qatar^32^. While immunological assessments for all vaccine trials are underway and correlates of protection are not yet determined, these data highlight the potential impact that reduced neutralization capacity to B.1.351 may have on protection against mild symptomatic COVID-19 across various platforms. Though comparable to BNT162b2 in other settings, human efficacy trials with mRNA-1273 have not been conducted in regions where B.1.351 circulates as a dominant variant.

Vaccine development for COVID-19 has benefitted from clinically translatable data from the NHP^33-39^. As there have been no published studies on vaccine protection in NHP challenged with the B.1.351 variant, we evaluated the impact of the dose and number of immunizations with mRNA-1273 on immunogenicity and protection against B.1.351 challenge in NHP.

## METHODS

### Pre-clinical mRNA-1273 mRNA and Lipid Nanoparticle Production Process

A sequence-optimized mRNA encoding prefusion-stabilized SARS-CoV-2 S-2P^40,41^ protein was synthesized *in vitro*. The mRNA was purified by oligo-dT affinity purification and encapsulated in a lipid nanoparticle through a modified ethanol-drop nanoprecipitation process described previously^42^.

### Rhesus Macaque Model

Animal experiments were carried out in compliance with US National Institutes of Health regulations and approval from the Animal Care and Use Committee of the Vaccine Research Center and Bioqual, Inc. (Rockville, MD). Challenge studies were conducted at Bioqual, Inc. Male and female, 3-12 year-old, Indian-origin rhesus macaques were sorted by sex, age and weight **(Supplemental Appendix)** and then stratified into groups of four. NHP were immunized intramuscularly (IM) at week 0 and week 4-5 with either 30 or 100 µg mRNA-1273 in 1 mL of 1X PBS into the right hind leg or 30 µg at week 0. Naïve aged-matched NHP were included as controls. At week 12 (7-8 weeks post-boost or 12 weeks after the single vaccination), all NHP were challenged with a total dose of 5×10^5^ PFU of SARS-CoV-2 B.1.351 strain. The viral inoculum was administered as 3.75×10^5^ PFU in 3 mL intratracheally (IT) and 1.25×10^5^ PFU in 1 mL intranasally (IN) in a volume of 0.5 mL into each nostril. Pre- and post-challenge sample collection is detailed in **Figure S1**.

### Quantification of SARS-CoV-2 RNA and sgRNA

At the time of collection, NS were frozen in 1 mL of 1X PBS containing 1 μL of SUPERase-In RNase Inhibitor (Invitrogen) and BAL was mixed with 1 mL of RNAzol BD containing 10 μL acetic acid and both were frozen at -80°C until extraction. Extraction and quantitation of sgRNA envelope (E) and nucleocapsid (N) were performed as previously described^38^.

### 10-plex Meso Scale ELISA

Multiplexed plates (96-well) precoated with SARS-CoV-2 S-2P^41^ and RBD proteins from the following strains: WA-1, B.1.351, B.1.1.7, and P.1., SARS-CoV-2 N protein, and Bovine Serum Albumin (BSA) are supplied by the manufacturer [Meso Scale Display (MSD)]. Determination of antibody binding was performed as previously described^43^.

### 4-plex Meso Scale ELISA

MSD SECTOR^®^ plates are precoated by the manufacturer with SARS-CoV-2 proteins (S-2P^41^, RBD, and N) and a BSA control in each well in a specific spot-designation for each antigen. Determination of antibody binding was performed as previously described^38^.

### Meso Scale ELISA for Mucosal Antibody Responses

Using previously described methods^44^, total S-specific IgG and IgA were determined by MULTI-ARRAY ELISA using Meso Scale technology (Meso Scale Discovery, MSD).

### Lentiviral Pseudovirus Neutralization Assay

As previously described^45^, pseudotyped lentiviral reporter viruses were produced by the co-transfection of plasmids encoding S proteins from Wuhan-1 strain (Genbank #: MN908947.3) with a D614G mutation, a luciferase reporter, lentivirus backbone, and human transmembrane protease serine 2 (TMPRSS2) genes into HEK293T/17 cells (ATCC CRL-11268). Similarly, pseudoviruses containing S from B.1.351, P.1, and B.1.1.7 were produced. Sera, in duplicate, were tested for neutralizing activity against the pseudoviruses by quantification of luciferase activity [in relative light units (RLU)].

### VSV Pseudovirus Neutralization Assay

To make SARS-CoV-2 pseudotyped recombinant VSV-ΔG-firefly luciferase virus, BHK21/WI-2 cells (Kerafast, EH1011) were transfected with the Wuhan-1 strain (Genbank #: MN908947.3) S plasmid expressing full-length S with D614G mutation or S of B.1.351. Neutralization assays were completed on A549-ACE2-TMPRSS2 cells with serially diluted serum samples as previously described^26^.

### Focus Reduction Neutralization Test (FRNT)

Viruses were propagated in Vero-TMPRSS2 cells to generate viral stocks. Viral titers were determined by focus-forming assay on VeroE6 cells. Viral stocks were stored at -80°C until use. FRNT assays were performed as previously described ^46^.

### Statistical Analysis

Graphs show data from individual NHP with dotted lines indicating assay limits of detection. Groups were compared by Kruskal-Wallis test, followed by pairwise Wilcoxon Rank-sum tests with Holm’s adjustment on the set of pairwise tests if the Kruskal-Wallis was significant, for the primary analysis of viral load at day 2 in the BAL and NS, as well as other comparisons between dose groups. Correlations were estimated and tested using Spearman’s nonparametric method. Linear regression was used to explore the relationship between antibody levels and sgRNA, including quadratic terms for comparing the newly generated data and that previously published (ref)^47^, with likelihood ratio tests to compare models and assess interaction effects.

## RESULTS

### Humoral and mucosal antibody responses following mRNA-1273 vaccination

Vaccination of NHP with 10-100 µg of mRNA-1273 at weeks 0 and 4 conferred rapid and complete control of detectable viral replication in both the upper and lower airways following SARS-CoV-2 USA/Washington-1 (WA-1) challenge^33,38^. In the current study, to assess the influence of dose and number of immunizations on immunogenicity and protection against B.1.351, NHP were immunized with 30 or 100 µg in the standard 0- and 4-week vaccine regimen (x2) or a single dose (x1) of 30 µg **(Figure S1)**.

We first performed a temporal analysis of serum neutralizing antibody responses after single immunization or prime and boost with mRNA-1273. Neutralizing responses of 50 and 99 geometric mean reciprocal ID_50_ titers (GMT) were detected against D614G by 2 weeks after a single immunization with 30 or 100 µg of mRNA-1273, respectively. Consistent with our prior study^38^, there was a ∼100-fold increase in D614G-specific neutralizing antibodies following a boost with 100 or 30 µg of mRNA-1273. All 8 of the 100 µg x2 immunized NHP and 7 of 8 receiving 30 µg x2 had >10^3^ reciprocal ID_50_ titers **(Figures 1E-G and S2A)** and 14/16 animals from the 2-dose regimen had detectable neutralizing activity against B.1.351 (**Figures 1E-G and S2B**). By contrast, only 6/8 NHP that received a single dose of 30 µg had detectable neutralizing responses against D614G **(Figures 1E-G and S2A)**, and none (0/8) had detectable neutralizing antibodies against B.1.351 **(Figures 1E-G and S2B)**. Following boost in the 30 and 100 µg x2 groups, neutralizing antibodies against B.1.351 remained >10^2^ and >10^3^ reciprocal ID_50_ GMT, respectively **(Figures 1E-G and S2B)**. These data support the importance of boosting to increase neutralizing responses against the B.1351 variant.

**Figure 1.**
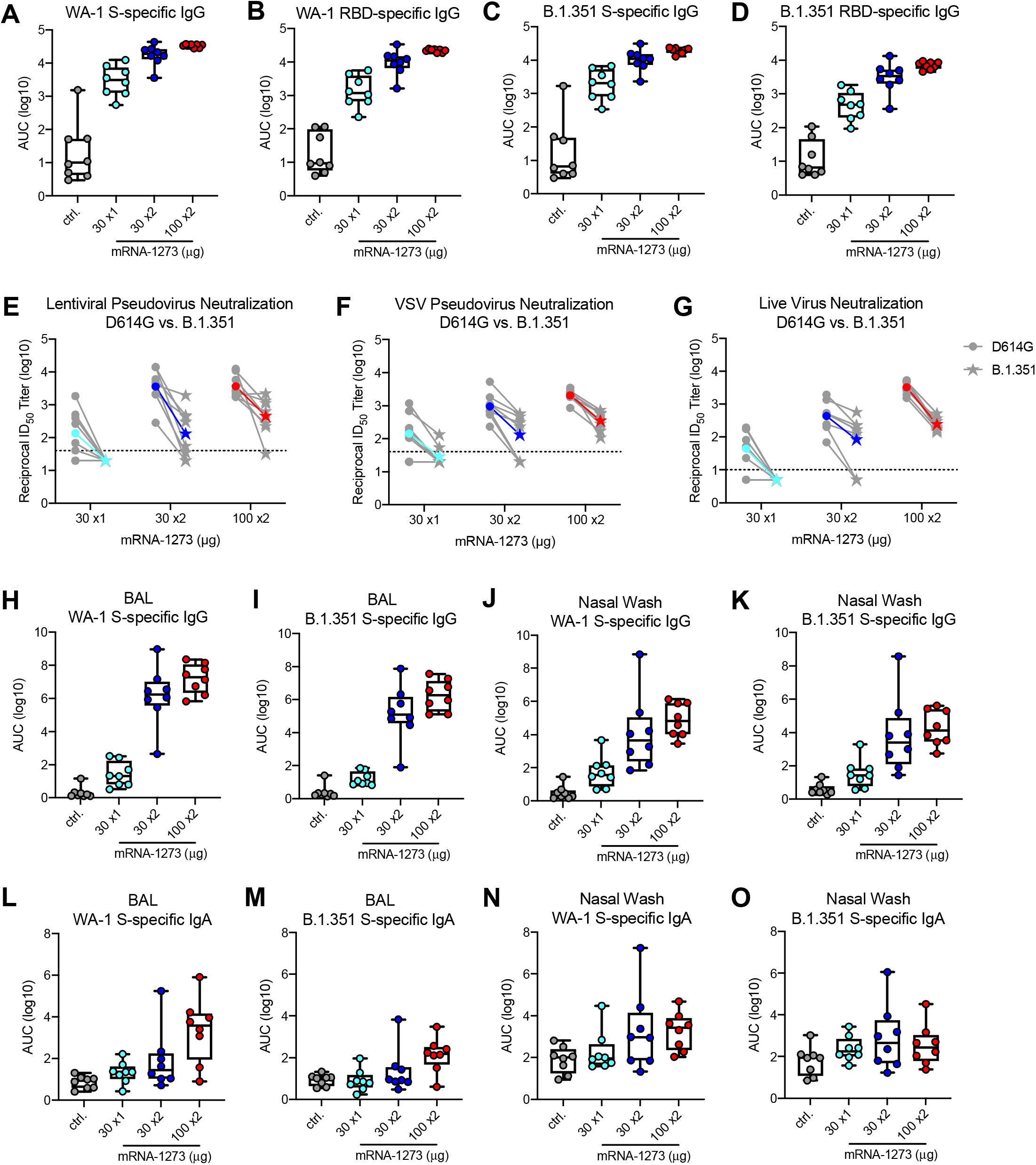
Antibody responses following mRNA-1273 immunization. Rhesus macaques were immunized with mRNA-1273 (30 µg, one dose – light blue; 30 µg, two doses – dark blue; or 100 µg, two doses – red), according to Figure S1. Aged-matched naïve NHP (gray) were used as controls. Sera collected at week 12, immediately before challenge, were assessed for SARS-CoV-2 USA/Washington1 (WA-1) (A-B) and B.1.351 (C-D) S-specific (A, C) and RBD-specific (B, D) IgG by MULTI-ARRAY ELISA, SARS-CoV-2 D614G and B.1.351 lentiviral-based pseudovirus neutralization (E), VSV-based pseudovirus neutralization (F), and focus reduction neutralization (G). BAL (H-I, L-M) and nasal washes (J-K, N-O) collected at week 7 were assessed for SARS-CoV-2 WA-1 (H, J, L, N) and B.1.351 (I, K, M, O) S-specific IgG (H-K) and IgA (L-O) by MULTI-ARRAY ELISA. (A-D, H-O) Circles represent individual NHP. Boxes and horizontal bars denote the IQR and medians, respectively; whisker end points are equal to the maximum and minimum values. (E-G) Gray lines represent individual NHP, and colored lines represent geometric mean titers (GMT). Dotted lines indicate neutralization assay limits of detection. Symbols represent individual NHP and may overlap for equal values.

Focusing on the time of challenge, ∼8 weeks post-boost or 12 weeks after the single immunization, S-specific binding and neutralizing antibody responses were assessed. Using a 10-plex MULTI-ARRAY ELISA, we assessed WA-1 and B.1.351 S- and RBD-specific antibody binding responses (**Figure 1A-D)**, which represent the vaccine and challenge strains, respectively. Binding to S and RBD proteins representing the P.1 lineage, which is prominent in Brazil, and the highly transmissible B.1.1.7 variant, which is circulating globally, were also assessed (**Figure S3A-D**). There was a vaccine dose-dependent increase in S- and RBD-specific antibody responses against WA-1 **(Figure 1A-B)** and B.1.351 **(Figure 1C-D)**, where, for example, a single dose of 30 µg of mRNA-1273 elicited 1,800 area under the curve (AUC) units for B.1.351 S-specific antibody responses, and two doses of 100 µg elicited 19,000 AUC. B.1.1.7 **(Figure S3A-B)**. P.1 **(Figure S3C-D)** S- and RBD-specific responses were similarly dose-dependent.

*In vitro* neutralizing activity was next determined using two orthogonal pseudovirus neutralization assays and a live virus assay. Neutralizing responses against B.1.351 was compared to D614G, the benchmark strain that is being used for correlates analysis of the Phase 3 studies with mRNA-1273 and other vaccines. Using a D614G lentiviral-based pseudovirus neutralization assay, the reciprocal ID_50_ GMT was ∼3,600 following two doses of 100 µg. Consistent with the 8-fold reduction reported by us and others using human vaccine or convalescent serum^26,43,48-50^, the reciprocal ID_50_ GMT against B.1.351 was ∼450. Notably, in NHP that received a single 30 µg dose of mRNA, the reciprocal ID_50_ GMT against D614G was ∼150, but there were no detectable neutralizing antibodies against B.1.351 **(Figure 1E)**. We observed similar outcomes using VSV-based pseudovirus **(Figure 1F)** and live virus **(Figure 1G)** neutralization assays. To extend the analysis, neutralization against the B.1.1.7 and P.1 variants was assessed. There was little change in neutralization in any vaccine group comparing D614G to B.1.1.7 **(Figure S3E)**; however, the reduction in neutralization against the P.1 variant compared to D614G was similar to that observed with B.1.351 **(Figure S3F)**. Taking all of these data together, antibody binding and neutralization responses were highly correlated with one another **(Figure S4)**.

To extend the antibody analyses to the mucosal sites of infection, S-specific IgG and IgA in BAL and nasal wash samples were assessed at ∼3 weeks post-boost or 7 weeks after the single 30 µg immunization. Consistent with systemic humoral responses, there was a dose-dependent increase in BAL and nasal wash WA-1 or B.1.351 S-specific IgG and IgA **(Figure 1H-O)**. BAL WA-1 **(Figure 1H)** or B.1.351 **(Figure 1I)** S-specific IgG titers following two doses of 30 or 100 µg of mRNA-1273 ranged from a GMT of 10^4^ - 10^7^ AUC, and nasal wash WA-1 **(Figure 1J)** or B.1.351 **(Figure 1K)** S-specific IgG titers ranged from 10^2^ – 10^5^AUC. For BAL, there was also a dose-dependent trend for WA-1 **(Figure 1L)** or B.1.351 **(Figure 1M)** S-specific IgA titers, albeit to lower magnitude than for IgG. mRNA-1273 did not induce notable upper airway WA-1 **(Figure 1N)** or B.1.351 **(Figure 1O)** S-specific IgA responses as nasal wash IgA levels in all vaccine groups were similar to control NHP. Overall, mRNA-1273 vaccination elicits WA-1 and B.1.351 S-specific IgG and IgA antibodies in serum and lower airways and IgG in the upper airways as previously shown^38^.

### T cell responses following mRNA-1273 vaccination

mRNA 1273 induces Th1, CD4 T follicular helper (Tfh) responses and CD8 T cells in NHP and humans^33,51,52^. Consistent with these data, S-specific Th1 responses were induced in a dose-dependent manner with higher responses in the 100 µg dose group **(Figure 2A)**. There were low to undetectable Th2 responses in all vaccine groups **(Figure 2B)**. There was also dose-dependence in the frequency of S-specific Tfh responses expressing the surface marker CD40L **(Figure 2C)** or the canonical cytokine IL-21 **(Figure 2D)**, which are critical for improving antibody responses. S-specific CD8 T cell responses were observed in 5/8 NHP that received two doses of 100 µg mRNA-1273 **(Figure 2E)**. These data show that mRNA-1273 induces Th1- and Tfh-skewed CD4 responses and CD8 T cells at the highest dose.

**Figure 2.**
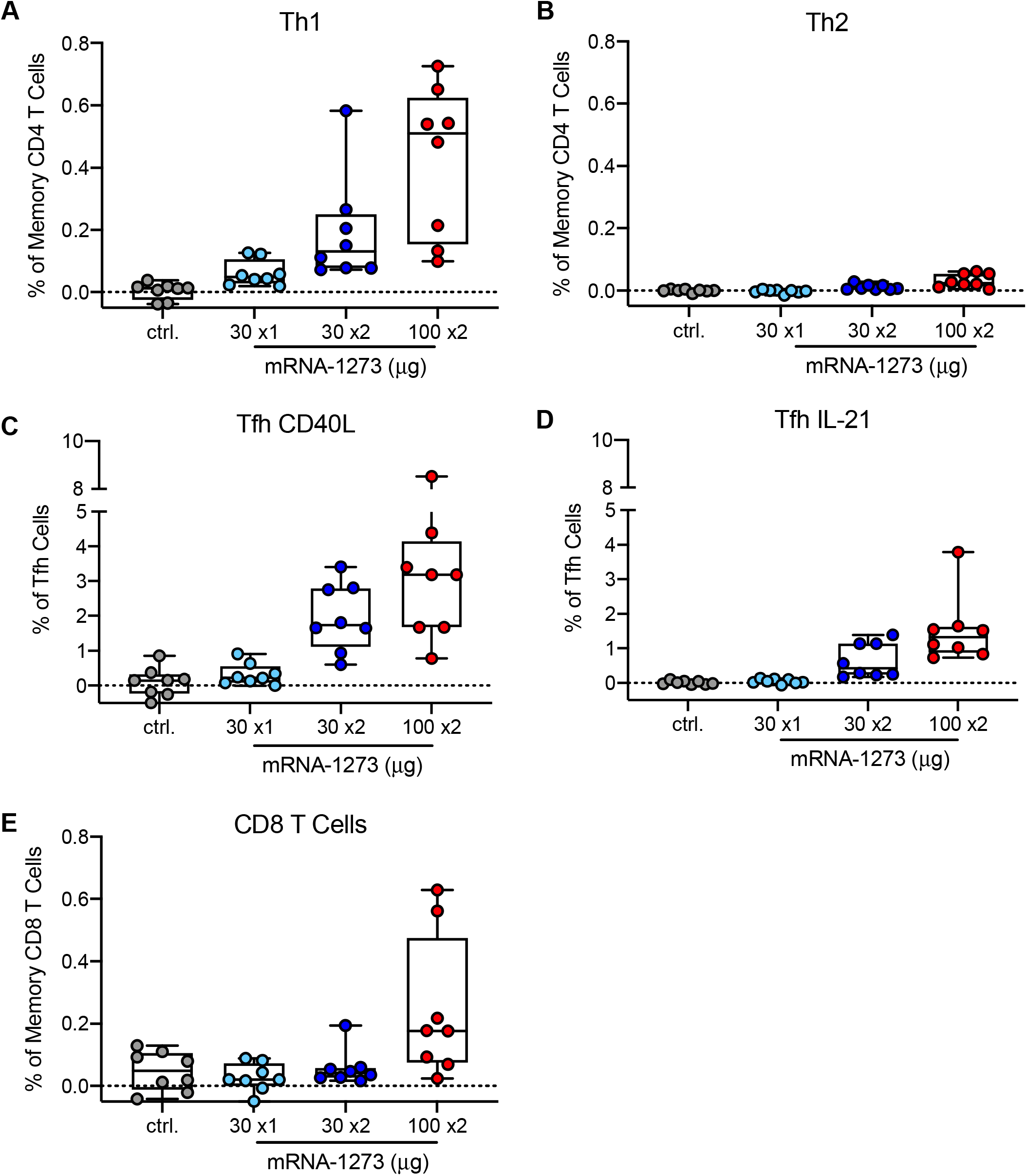
T cell responses following mRNA-1273 immunization. Rhesus macaques were immunized according to Figure S1. Intracellular staining was performed on PBMCs at week 7 to assess T cell responses to SARS-CoV-2 S protein peptide pools, S1 and S2. Responses to S1 and S2 individual peptide pools were summed. (A) Th1 responses (IFNg, IL-2, or TNF), (B) Th2 responses (IL-4 or IL-13), (C) Tfh CD40L upregulation (peripheral follicular helper T cells (Tfh) were gated on central memory CXCR5^+^PD-1^+^ICOS^+^ CD4 T cells), (D) Tfh IL-21, (E) CD8 T cells. Boxes and horizontal bars denote IQR and medians, respectively; whisker end points are equal to the maximum and minimum values. Circles represent individual NHP. Dotted lines are set to 0%.

### Protective Efficacy against SARS-CoV-2 Replication in Upper and Lower Airways

As this is the first study to challenge NHP with B.1.351, an extensive analysis was performed to characterize the sequence and *in vivo* pathogenicity. Deep sequencing was performed after each passage (P) of the B.1.351 strain, which was first isolated at Johns Hopkins University (JHU). P1 and P2 stocks were compared to the B.1.351 reference strain and the original JHU clinical isolate and retained all mutations in the RBD and NTD sites on the spike protein **(Figure S5A)**. The JHU B.1.351 P2 stock was then administered to Golden Syrian hamsters, a highly pathogenic SARS-CoV-2 animal model, at three different concentrations to characterize weight loss **(Figure S5B)** and in NHP to measure upper and lower airway viral replication by qRT-PCR for sgRNA **(Figures S5C-D)**. Based on these data, a B.1.351 challenge dose of 5×10^5^ PFU was selected for the vaccine study; this dose would induce sgRNA levels similar to the higher values obtained from nasal secretions of humans following SARS-CoV-2 infection^53,54^.

To assess protective efficacy of mRNA-1273 against the B.1.351 SARS-CoV-2 variant, the NHP were challenged with a total dose of 5×10^5^ PFU of B.1.351 by intratracheal (IT) and intranasal (IN) routes 7-8 weeks post-boost for the two-dose regimens and 12 weeks after the single dose regimen **(Figure S1)**. Two days post-challenge, only 2 of 8 NHP that received 100 µg of mRNA-1273 had detectable SARS-CoV-2 envelope (E) sgRNA (sgRNA _E) in BAL compared to 8 of 8 in the control group **(Figure 3A)**. By sgRNA_E **(Figure 3A)** or nucleocapsid (N) sgRNA (sgRNA _N) qRT-PCR **(Figure 3C)**, the 100 µg group had a significant decrease in viral load compared to NHP that received a single dose of 30 µg (*p* = 0.0054) or to the control NHP (*p* = 0.0009). NHP that received 30 µg x2 also had a significant decrease in viral load compared to control NHP (*p* = 0.0054) and showed a trend toward reduced viral replication compared to a single immunization with 30 µg. NHP that received a single vaccination with 30 µg showed a trend toward reduced viral replication compared to control NHP. At day 4 post-challenge, the pattern was the same, with significant reduction in viral replication in BAL for all vaccine groups compared to control NHP, and for the 100 µg group compared to 30 µg x1 **(Figures 3A,C)**. By day 7, while 7 of 8 control NHP still had ∼4 logs of sgRNA_E, there was no detectable sgRNA_E in 6 of 8 NHP in all vaccine groups **(Figure 3A)**, consistent with control of viral replication in the lower airway. In addition, the inability to culture virus from the BAL of 4/8 and 7/8 NHP immunized with 30 and 100 µg x2 of mRNA-1273, respectively, two days post-challenge further confirms the ability of mRNA-1273 to control lower airway viral replication **(Figure 3E)**. Moreover, on day 2 post-challenge, BAL viral titers and sgRNA were highly correlated **(Figures 3F-G)**, where no virus was culturable from BAL of all NHP with BAL sgRNA_N <1.2×10^4^ RNA copies/mL **(Figure 3G)**.

**Figure 3.**
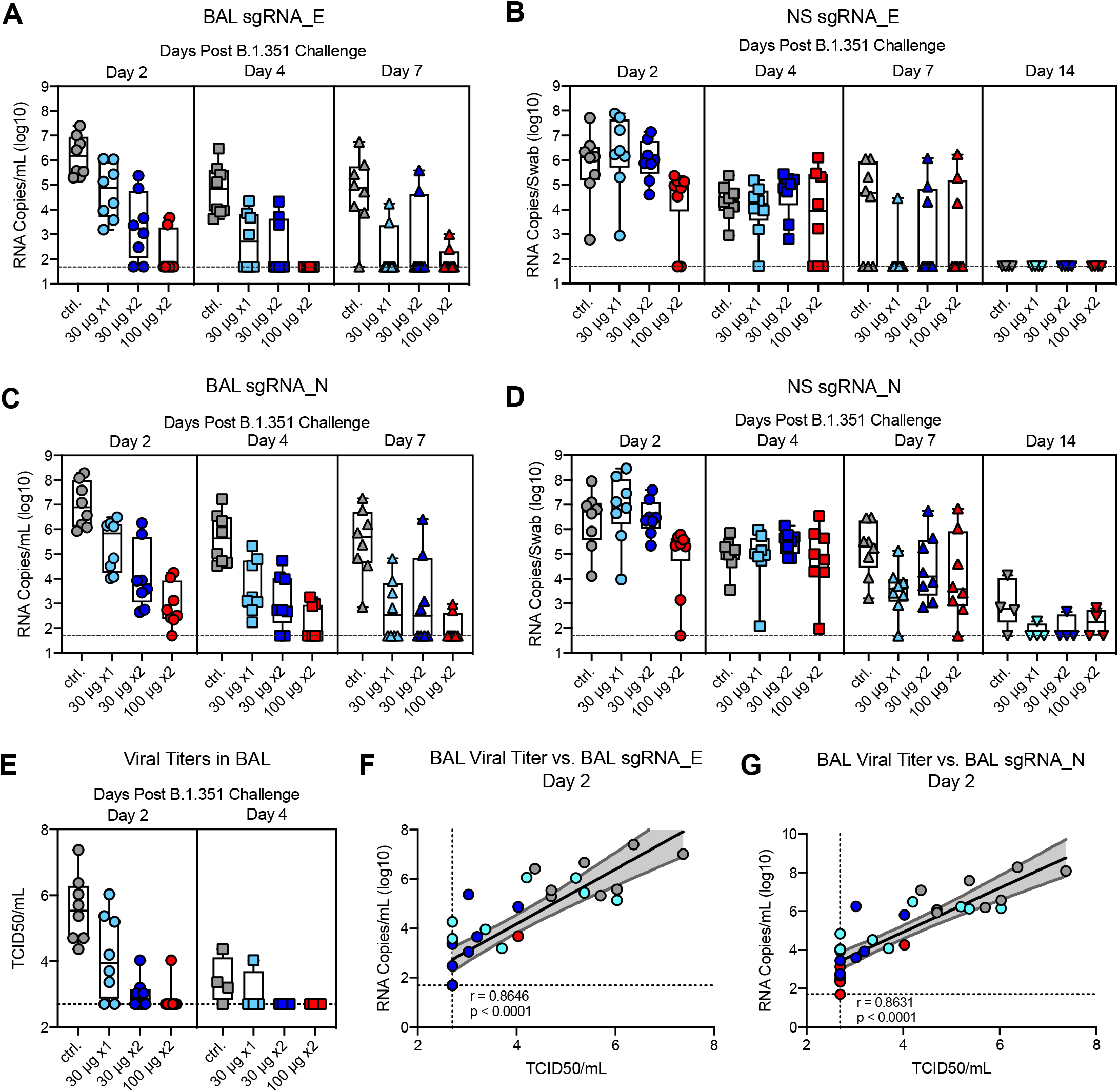
Efficacy of mRNA-1273 against upper and lower respiratory B.1.351 viral replication. Rhesus macaques were immunized and challenged as described in Figure S1. BAL (A, C) and nasal swabs (NS) (B, D) were collected on days 2 (circles), 4 (squares), and 7 (triangles), and 14 (inverted triangles) post-challenge, where applicable, and viral replication was assessed by detection of SARS-CoV-2 E-(A-B) and N-specific (C-D) sgRNA. (E) Viral titers were assessed by TCID50 assay for BAL collected on days 2 and 4 post-challenge. Boxes and horizontal bars denote the IQR and medians, respectively; whisker end points are equal to the maximum and minimum values. (F-G) Plots show correlations between viral titers and sgRNA_E (F) and sgRNA_N (G) in BAL 2 days post-challenge. Black and gray lines indicate linear regression and 95% confidence interval, respectively. ‘r’ and ‘p’ represent Spearman’s correlation coefficients and corresponding p-values, respectively. Symbols represent individual NHP and may overlap for equal values.

In contrast to the significant reduction of viral replication in the lower airway, at day 2 post-challenge, the only significant reduction in viral replication for sgRNA_E **(Figure 3B)** or sgRNA _N **(Figure 3D)** in nasal swabs (NS) was in the 100 µg dose group. The differences in NHP that received 100 µg x2 were marginally lower compared to the groups that received 30 µg x2 or 30 µg x1 (*p* = 0.0273 and 0.0350, respectively) with no other significant pairwise differences. The groups were not significantly different at the later time points. A notable finding was that 5 of 8 control NHP still had ∼4 log_10_ of sgRNA_E and all 8 NHP had sgRNA_N at day 7 in the NS, highlighting persistence of sgRNA following B.1.351 through day 7 post-challenge. With that, there was ∼4 log_10_ of sgRNA_E in the NS of 3 of 8 NHP in the 100 µg x2 group. Overall, these data show significant reduction and rapid control of B.1.351 viral replication in the lower airways of mRNA-1273 immunized NHP with more limited control in the upper airway.

### Inflammation and viral load in lung tissue post-challenge

To provide a further assessment of protection following vaccination, NHP in each of the dose groups were assessed for virus-related pathology and for the detection of viral antigen (VAg) in the lung 8 days post B.1.351 challenge. The severity of inflammation, which ranged from minimal to moderate, was similar across lung samples from NHP that received vaccine in doses of 100 µg x2, 30 µg x2, or 30 µg x1 **(Figure S6)**. The inflammatory lesions in the lung were characterized by a mixture of lymphocytes, histiocytes and fewer polymorphonuclear cells associated with variably expanded alveolar capillaries, occasional areas of perivascular inflammation, and Type II pneumocyte hyperplasia. Two out of 4 NHP that received 30 µg x1 of mRNA-1273 vaccine had trace amounts of virus detected in the lung. There was no detection of VAg in any lung sample from NHP that received two doses of 30 or 100 µg. All 4 NHP in the control group had variable amounts of VAg detected in the lung **(Figure S6, Table S1)**.

### Post-challenge humoral and mucosal antibody responses

The assessment of antibody responses post-challenge has been useful for determining whether viral replication in the BAL or NS is sufficient to boost vaccine-induced anamnestic S-specific antibody responses in these mucosal tissues^38,44^. In BAL, WA-1 and B.1351 S-specific IgG **(Figures S7A-B)** or IgA **(Figures S7C-D)** responses did not increase post-challenge in NHP that received two immunizations of 30 or 100 µg of mRNA-1273. However, by 14 days post-challenge, there was an increase in WA-1 and B.1.351 S-specific IgG responses in NHP that received 30 µg x1 of mRNA-1273, to levels that were similar to the NHP that received 30 or 100 µg x2 and higher than the unvaccinated controls **(Figures S7A-D)**. In NS, there was an increase in WA-1 and B.1.351 S-specific IgG **(Figures S7E-F)** or IgA **(Figures S7G-H)** responses in unvaccinated NHP and those immunized with 30 µg of mRNA-1273 once or twice; however, there were no anamnestic S-specific antibody responses in 100 µg dose group. Overall, these data show that the increase in S-specific anamnestic antibody responses in both the BAL and NS are associated with viral replication in these mucosal sites and may explain the relatively rapid clearance of virus from NS in the 30 µg x1 dose group.

### Antibody correlates of protection

Assessing immune correlates of protection following vaccination is a critical aspect of vaccine development. We recently reported that mRNA-1273 induced antibody responses are a mechanistic correlate for reducing viral replication against WA-1 challenge in NHP^38^. Here, B.1.351 S-specific IgG antibody titers at week 12, the time of challenge, also correlated strongly with reduction of sgRNA in both BAL **(Figure 4A)** and NS **(Figure 4D)** at day 2 post-challenge. In addition, both pseudovirus and live viral neutralization correlate significantly with reduction of sgRNA in both BAL **(Figure 4B-C)** and NS **(Figures 4E-F)**.

**Figure 4.**
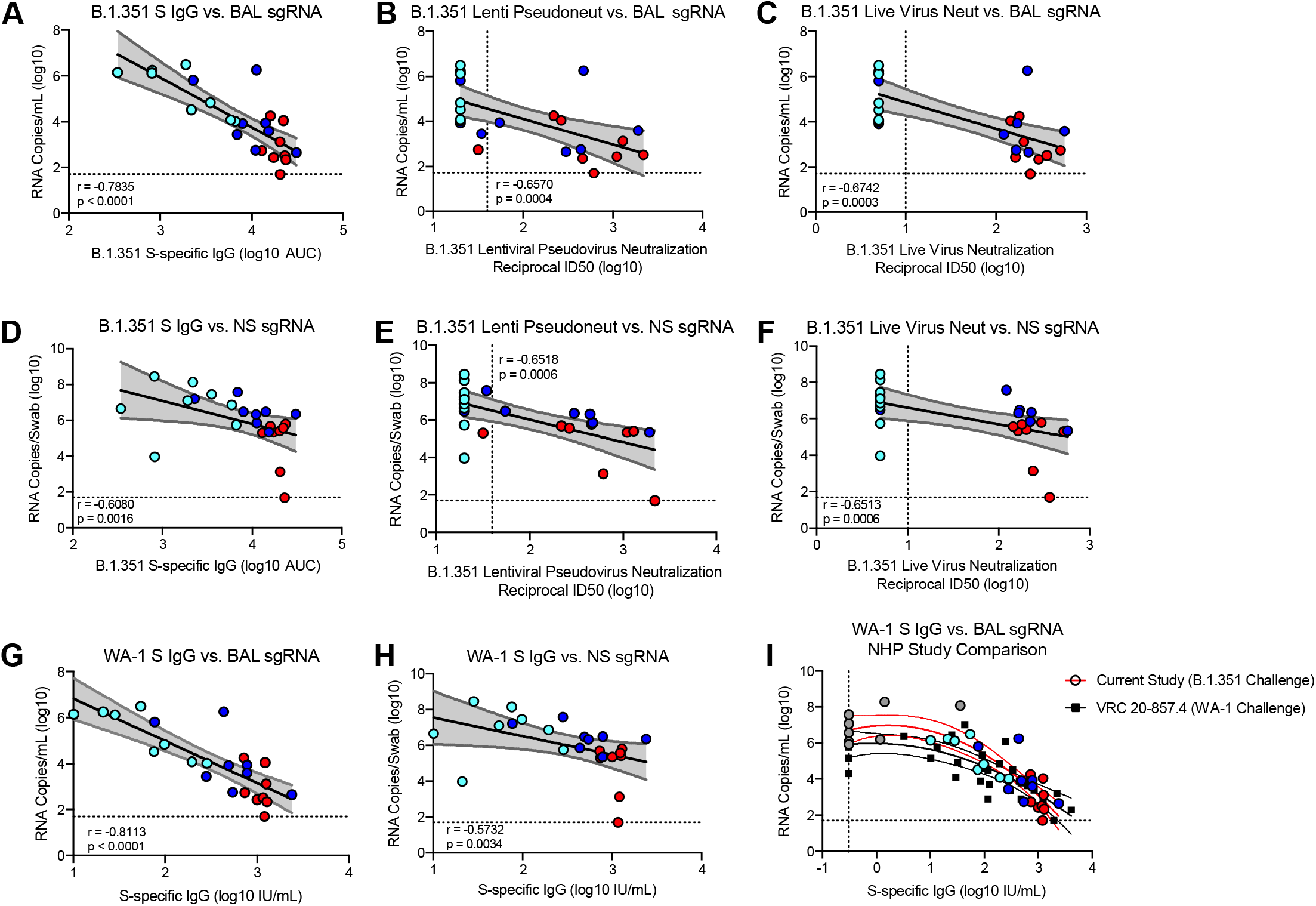
Antibody correlates of protection. Rhesus macaques were immunized and challenged as described in Figure S1. (A-F) Plots show correlations between week 12 SARS-CoV-2 B.1.351 S-specific IgG (A, D), lentiviral-based pseudovirus neutralization (B, E), and focus reduction neutralization (C, F) with N-specific sgRNA in BAL (A-C) and NS (D-F) at day 2 post-challenge. (G-H) Plots show correlations between week 12 SARS-CoV-2 WA-1 S-specific IgG, converted to IU/mL, with N-specific sgRNA in BAL (G) and NS (H) at day 2 post-challenge. Circles represent individual NHP, where colors indicate mRNA-1273 dose. Dotted lines indicate assay limits of detection. (I) The relationship between pre-challenge WA-1 S-specific IgG and day 2 BAL sgRNA_N, with data from the current study using a B.1.351 challenge (filled circles, red curve fit) superimposed on data from our previous study with a WA-1 challenge (black squares, black curve fit); lines indicate quadratic curve fit and 95% confidence intervals. Symbols represent individual NHP and may overlap for equal values.

In a recent report on correlates of protection following mRNA-1273 immunization in NHP, we established a linear relationship between WA-1 S-specific antibody titers as defined by international units (IU) and subsequent sgRNA 2 days after SARS-CoV-2 WA-1 challenge. Here, we similarly converted pre-challenge WA-1 S-specific antibody titers to IU **(Table S2)**. Again, pre-challenge WA-1 S-specific IgG titers and sgRNA in BAL **(Figure 4G)** and NS **(Figure 4H)** at day 2 post-challenge were negatively correlated. Based on a linear model, vaccinated animals that had S-specific IgG of 100 and 120 IU/mL were predicted to have sgRNA_N and sgRNA_E, respectively, in BAL of approximately 2 log_10_ lower than the average for PBS controls. Serum neutralizing activity was associated with control of viral replication; 13/14 of the NHP with detectable pseudovirus neutralizing activity against B.1.351 had BAL sgRNA_N <10^5^ **(Figure 4B)**. A similar linear relationship between S-specific antibody levels and viral replication in nasal swabs was apparent, although few of the NHP had NS sgRNA_N below 10^5^.

As the antibody levels of S-specific IgG are very similar to those we had previously reported using NHP challenged with the SARS-CoV-2 WA-1^38^, we conducted an additional exploratory analysis of the data from both studies to investigate the consistency of the relationship between S-specific antibodies and BAL sgRNA, across the two different viruses. For low S-specific IgG levels, sgRNA was higher for animals challenged with B.1.351, but for animals with levels of S-specific IgG greater than approximately 100 IU/mL, the estimated regression curve slopes are similar as are the levels that correspond to 2-4 log_10_ reductions in sgRNA in BAL compared to controls **(Figure 4I)**. These data suggest a similar relationship between S-specific IgG levels and lower airway protection against homologous and variant SARS-CoV-2 strains in NHP.

## DISCUSSION

mRNA-1273 and BNT162b2 vaccines were shown to be ∼95% effective in clinical trials performed in the US, during which times WA-1 and D614G variants circulated most widely^2,55^. A critical issue is whether these and other vaccines will mediate protection against the rapidly emerging variants. At present, the B.1.351 variant is one of greatest concern compared to WA-1, D614G, or B.1.1.7 based on the higher reduction in neutralization using vaccine sera^7-17^ and clinical trials showing lower efficacy against mild infection^29-31^. Here, we present evidence that mRNA-1273 can significantly reduce viral replication in the upper and lower airways and prevent or limit inflammation in the lung following B.1.351 challenge in NHP. Importantly, the dose of the vaccine and number of immunizations had a significant effect on the protective capacity. These data are consistent with a recent report showing that vaccine effectiveness against PCR-confirmed infection with the B.1.351 variant is 16% after 1 dose of BNT162b2 and 75% after two doses^32^.

Antibodies play a critical role in mediating vaccine-elicited protection against SARS-CoV-2 in NHP models^33,37,39,44,56,57^. Here, we show that there was a dose-dependent increase in WA-1 and B.1.351 S- and RBD-specific antibody titers and D614G and B.1.351 neutralization titers **(Figure 1A-D)**. Of note, there was ∼5-10-fold reduction in sensitivity to neutralization for B.1.351 compared to D614G, consistent with many reports assessing responses from humans that received mRNA 1273 or other vaccines. There was also a dose-dependent increase in WA-1 and B.1351 S-specific IgG in BAL and nasal washes, where IgA was detected only in the 100 µg group. Importantly, while there were detectable neutralization titers against D614G following a single immunization with 30 µg, these NHP had no detectable neutralization against B.1.351. By contrast, neutralizing titers against B.1.351 following a second immunization with 30 or 100 µg increased to ∼10^3^ reciprocal ID_50_ titer. These data highlight the importance of a prime and boost regimen for optimizing neutralization antibody responses, particularly against B.1.351 and likely for any other variant of concern for which vaccine-induced neutralization is decreased. Last, the frequency of S-specific Th1 and Tfh responses were also dose-dependent, with CD8 T cell responses detected in blood only in NHP receiving the 100 µg dose. These data corroborate previous studies by us and others that have shown a direct correlation between CD4 T cell responses, most notably Tfh cells, and improved magnitude and function of antibody responses in NHP^33,38,51,58^.

As this was the first study to use the B.1.351 variant for challenge in NHP, extensive sequence analyses were performed to propagate a challenge stock with a matched S sequence as compared to the reference isolate^59^. Naïve NHP infected with the B.1.351 stock notably had peak sgRNA levels of ∼10^7^ copies/mL in BAL, which is higher than reported by us and others for challenge studies using the WA-1 strain^33,37,39,44,56,57^. At 7 days post-challenge, most of the control NHP still had ∼10^5^ RNA copies/mL and copies/swab present in BAL and NS, respectively. This contrasts with the more rapid and complete reduction of viral replication following challenge with the WA-1 strain^33,37,39,44,56,57^ and higher than viral load typically seen in human infections. Whether the amount and persistence of the B.1351 virus *in vivo* in this study relates to the challenge dose or suggests that this variant has inherent properties that make it more difficult to control and clear compared to the WA-1 strain is a focus of ongoing analyses.

The dose of B.1351 used here provided a stringent challenge for vaccine-elicited protection. Nonetheless, there was a significant reduction in viral replication in BAL at day 2 post-challenge in the groups that received 30 and 100 µg twice; 6 of 8 NHP that received the clinically relevant 100 µg vaccine regimen had no detectable sgRNA_E in BAL. Of note, while there was no detectable serum neutralizing activity against B.1351 virus with a single immunization of 30 µg, there was still significant reduction in viral replication in BAL by day 4 post-challenge compared to controls with limited inflammation or viral antigen in the lungs at day 7. These data are consistent with our recent study in which NHP immunized with only 1 or 3 µg of mRNA-1273 twice demonstrated a reduction in viral replication in BAL and limited lung pathology following WA-1 challenge despite absence of detectable neutralizing activity^38^. There are several potential mechanisms for lower airway protection in the absence of detectable serum neutralizing antibodies. First, binding antibodies could mediate viral reduction through various Fc effector functions in the BAL as has been suggested as a mechanism for protection from vaccine-induced polyclonal serum^60,61^ and certain monoclonal antibodies^62,63^. Second, vaccine-primed anamnestic responses in the BAL in the first week post-challenge may enhance control of infection^38,44^. Third, current *in vitro* assays may not be sensitive enough to detect the full spectrum of *in vivo* neutralization activity, that may be present *in vivo*. Last, while it is possible that T cells are contributing to control of viral replication in the lower airway, there is no current evidence for this following vaccination or primary infection^64^ in the NHP model.

In contrast to protection observed in the lower airway against the B.1.351 challenge, there was more limited control of viral replication in the upper airway except in NHP that received 2 doses of 100 µg. These data are consistent with prior studies in NHP showing that a higher amount of antibody is required for reduction of viral replication in the upper airway than the lower airway following mRNA-1273 vaccination^33,38^. Moreover, recent results from human vaccine trials show that there is greater protection against severe disease than mild disease against the B.1.351 variant following immunization with Ad26.CoV2^29^ or BNT162b2^32^.

Serum antibody levels were strong predictors of reduction of viral load in BAL and NS. These data are consistent with studies by us and others showing that antibodies can be a mechanistic correlate of protection in NHP following vaccination^38^ or infection^64^. The correlation is most robust between S-specific IgG and BAL sgRNA; our modeling suggests that animals with S-specific IgG levels of approximately 100 IU/mL display sgRNA levels in BAL of approximately 2 log_10_ lower than the control animals, with a further decrease of approximately 2 log_10_ sgRNA for every 1 log_10_ increase in IgG IU/mL. We also noted that 13/14 animals with any detectable neutralizing antibodies against B.1.351 had BAL sgRNA_N <10^5^. Note that no virus was culturable from BAL of any NHP with BAL sgRNA_N <1.2×10^4^ RNA copies/mL. These data suggest that even for vaccines that elicit low to undetectable B.1.351-specific neutralizing antibodies, there can still be control of viral replication in the lower airway.

The NHP model has been critical for guiding vaccine development against COVID-19 in humans. This report provides evidence that the current two-dose regimen with mRNA-1273 is important for inducing higher neutralizing antibody responses, CD4 and CD8 T cell immunity and protection from viral replication in the lower and upper airways. The durability of immune responses and protection will likely be related to maintaining a combination of serum antibody and memory B cells that can rapidly respond to a new virus exposure. Moreover, immunity against B.1.351 or other variants of concern can be assessed by comparing reduction in neutralizing activity relative to D614G over time^43^. Ongoing studies are assessing how additional boosting with spike antigens based on WA-1 or variant strains will influence immunity and protection against B.1351 and other emerging variants in this model and in humans.

## Acknowledgements

We thank any additional members of all included laboratories for critical discussions and advice pertaining to experiments included in the manuscript. We thank Judy Stein and Monique Young for technology transfer and administrative support, respectively. We thank members of the NIH NIAID VRC Translational Research Program, including Chris Case, Hana Bao, Elizabeth McCarthy, Jay Noor, Alida Taylor, and Ruth Woodward, for technical and administrative assistance with animal experiments. We thank Huihui Mu and Michael Farzan for the ACE2-overexpressing 293 cells. We thank Adrian Creanga and Masaru Kanekiyo for the Vero-TMPRSS2 cells. We thank the laboratory of Peter Kwong for providing protein for use in ELISA assays for detection of mucosal antibodies. We thank Andy Pekosz for the B.1.351 variant. We thank Michael Brunner and Michael Whitt for kind support on recombinant VSV-based SARS-CoV-2 pseudovirus production. This work was supported by the Intramural Research Program of the VRC, NIAID, NIH. mRNA-1273 has been funded in part with Federal funds from the Department of Health and Human Services, Office of the Assistant Secretary for Preparedness and Response, Biomedical Advanced Research and Development Authority, under Contract 75A50120C00034. K.S.C.’s research fellowship was partially funded by the Undergraduate Scholarship Program, Office of Intramural Training and Education, Office of the Director, NIH. Virus propagation and live virus neutralization assays were funded by Emory Executive Vice President for Health Affairs Synergy Fund Award, Pediatric Research Alliance Center for Childhood Infections and Vaccines and Children’s Healthcare of Atlanta, and Woodruff Health Sciences Center 2020 COVID-19 CURE Award.

## Author Contributions

K.S.C., A.P.W., S.O., M.G., L.L., J.I.M., B.F., A.C., M.K, K.E.F., S.F.A., D.R.F., E. L., S.T.N., S.J.P., K.W.B., M.M., B.M.N., A.V.R., Z.F., T.S.J., E.B.M., P.M., A.R.H., F.L., B.C., M.P.,J.W., J.M.T., B.B., A.C., A.D., L.P., K.S., S.E., H.A., K.W., D.K.E., S.K., M.G.L., E.B., I.N.M., A.C., M.S.S., A.M., M.R., M.C.N., N.J.S., D.C.D., B.S.G., and R.A.S. designed, completed, and/or analyzed experiments. S.B-B. provided critical published reagents/analytic tools. K.S.C., M.C.N., and R.A.S wrote the manuscript. K.S.C., G.A., and M.C.N. prepared figures and tables. All authors contributed to discussions in regard to and editing of the manuscript.

