## Supplementary Appendix for "Evaluation of mRNA-1273 against SARS-CoV-2 B.1.351 Infection in Nonhuman Primates"

#### ADDITIONAL METHODOLOGICAL DETAILS

##### Quantification of SARS-CoV-2 RNA and sgRNA

At the time of collection, NS were frozen in 1 mL of 1X PBS containing 1 µL of SUPERase-In RNase Inhibitor (Invitrogen) and frozen at -80°C until extraction. Nasal specimens were thawed at 55°C, and the swab removed. The remaining PBS was mixed with 2 mL of RNAzol BD (Molecular Research Center) and 20 µL acetic acid. At the time of collection, 1 mL of BAL fluid was mixed with 1 mL of RNAzol BD containing 10 µL acetic acid and frozen at -80°C until extraction. BAL specimens were thawed at room temperature (RT) and mixed with an additional 1 mL of RNAzol BD containing 10 µL acetic acid. Total RNA was extracted from nasal specimens and BAL fluid using RNAzol BD Column Kits and eluted in 65 µL water. Subgenomic SARS-CoV-2 E and N mRNA was quantified via reverse transcription-polymerase chain reaction (PCR) as previously described (*cite Correlates paper here*). Reactions were conducted with 5 µL RNA and TaqMan Fast Virus 1-Step Master Mix (Applied Biosystems) with 500 nM primers and 200 nM probes. Primers and probes were as follows: sgLeadSARSCoV2_F: 5'-CGATCTCTTGTAGATCTGTTCTC-3', E subgenomic mRNA - E_Sarbeco_P: 5'-FAM-ACACTAGCCATCCTTACTGCGCTTCG-BHQ1-3' and E_Sarbeco_R: 5'-ATATTGCAGCAGTACGCACACA-3', N subgenomic mRNA - wtN_P: 5'-FAM-TAACCAGAATGGAGAACGCAGTGGG-BHQ1-3'and wtN_R: 5'-GGTGAACCAAGACGCAGTAT-3'. Reactions were run on a QuantStudio 6 Pro Real-Time PCR System (Applied Biosystems) at the following conditions: 50°C for 5 min, 95°C for 20 sec, and 40 cycles of 95°C for 15 sec and 60°C for 1 min. Absolute quantification was performed in comparison to a standard curve. For the standard curve, the E or N subgenomic mRNA sequence was inserted into a pcDNA3.1 vector (Genscript) and transcribed using MEGAscript T7 Transcription Kit (Invitrogen) followed by MEGAclear Transcription Clean-Up Kit (Invitrogen). The lower limit of quantification was 50 copies.

##### 10-plex Meso Scale ELISA

On the day of the assay, the plate is blocked for 60 minutes with MSD Blocker A (5% BSA). The blocking solution is washed off and test samples are applied to the wells at 4 dilutions (1:100, 1:500, 1:2500 and 1:10,000) unless otherwise specified and allowed to incubate with shaking for two hours. Plates are washed and SULFO-TAG™ labeled anti-IgG antibody is applied to the wells and allowed to associate with complexed coated antigen – sample antibody within the assay wells. Plates are washed to remove unbound detection antibody. A read solution containing ECL substrate is applied to the wells, and the plate is entered into the MSD Sector instrument. A current is applied to the plate and areas of well surface where sample antibody has complexed with coated antigen and labeled reporter will emit light in the presence of the ECL substrate. The MSD Sector instrument quantitates the amount of light emitted and reports this ECL unit response as a result for each sample and standard of the plate. Magnitude of ECL response is directly proportional to the extent of binding antibody in the test article. All calculations are performed within Excel and the GraphPad Prism software, v7.0. Readouts are provided as Area Under Curve (AUC).

**4-plex Meso Scale ELISA**

The assay will be performed with a Beckman Coulter Biomek based automation integration platform including the Biotek 405TS Plate Washer. Serum samples will be heat-inactivated for 30 min at 56ºC prior to assay. Plates are blocked for 60 min at RT with MSD blocker A solution without shaking. Plates are washed and MSD reference standard (calibrator), QC test sample (pool of COVID-19 convalescent sera) and human serum test samples are added to the precoated wells in duplicates in an 8-point dilution series. Reference standard is added in triplicates. MSD control sera (low, medium and high) are added undiluted in triplicates as per validated assay format. Additional assay controls might be added in triplicates. Samples are incubated at RT for 4 hr with shaking on a Titramax Plate shaker (Heidolph) at 1500 rpm. SARS-CoV-2 specific antibodies present in the sera or controls bind to the coated antigens. Plates are washed to remove unbound antibodies. Antibodies bound to the SARS-CoV-2 viral proteins are detected using an MSD SULFO-TAG™ anti-human IgG detection antibody incubated for 60 min at RT and with shaking. Plates are washed and a read solution (MSD GOLD™

read buffer) containing electrochemiluminescence (ECL) substrate is applied to the wells, and the plate is entered into the MSD MESO Sector S 600 detection system. An electric current is applied to the plates and areas of well surface which form antigen-anti human IgG antibody SULFO-TAG™ complex will emit light in the presence of the ECL substrate.

The MSD MESO Sector S 600 detection system quantitates the amount of light emitted and reports the ECL unit response as a result for each test sample, control sample and reference standard of each plate. Analysis is performed with the MSD Discovery Workbench software, Version 4.0. Calculated ECLIA parameters to measure binding antibody activities will include interpolated concentrations or assigned arbitrary units (AU/mL) read from the standard curve.

**Conversion to 4-plex Readout to International Units**

Recently the arbitrary units were bridged to the WHO International Standard [provided in international units/mL (IU/mL)], and a conversion factor was calculated and confirmed. Parallelism was established for all three antigens (SARS-CoV S-2P, RBD, and N) between the MSD provided reference standard and the WHO provided international standard. Concentration assignments were performed and then confirmed both at MSD and as part of a multi-site confirmation study.

**Lentiviral Pseudovirus Neutralization Assay**

Sera, in duplicate, were tested for neutralizing activity against the pseudoviruses by quantification of luciferase activity [in relative light units (RLU)]. Percent neutralization was normalized considering uninfected cells as 100% neutralization and cells infected with pseudovirus alone as 0% neutralization. $IC_{50}$ titers were determined using a log(agonist) vs. normalized-response (variable slope) nonlinear regression model in Prism v9.0.2 (GraphPad). For samples that do not neutralize at the limit of detection at 50%, a value of 20 was plotted and used for geometric mean calculations.

**VSV Pseudovirus Neutralization Assay**

which includes the following amino acid changes relative to that of Wuhan-Hu-1 (L18F, D80A, D215G, ΔLAL242-244, R246I, K417N, E484K, N501Y, D614G, A701V) and subsequently infected with VSVΔG-firefly-luciferase as previously described[1]. For samples that do not neutralize at the limit of detection at 50%, a value of 20 was plotted and used for geometric mean calculations.

**Focus Reduction Neutralization Test (FRNT)**

Briefly, samples were diluted at 3-fold in 8 serial dilutions using DMEM (VWR, #45000-304) in duplicates with an initial dilution of 1:10 in a total volume of 60 ml. Serially diluted samples were incubated with an equal volume of SARS-CoV-2 (100-200 foci per well) at 37° C for 1 hr in a round-bottomed 96-well culture plate. The antibody-virus mixture was then added to Vero cells and incubated at 37° C for 1 hr. Post-incubation, the antibody-virus mixture was removed and 100 µL of prewarmed 0.85% methylcellulose (Sigma-Aldrich, #M0512-250G) overlay was added to each well. Plates were incubated at 37° C for 24 hr. After 24 hr, methylcellulose overlay was removed, and cells were washed three times with PBS. Cells were then fixed with 2% paraformaldehyde in PBS (Electron Microscopy Sciences) for 30 min. Following fixation, plates were washed twice with PBS and 100 µl of permeabilization buffer (0.1% BSA [VWR, #0332], Saponin [Sigma, 47036-250G-F] in PBS), was added to the fixed Vero cells for 20 min. Cells were incubated with an anti-SARS-CoV S primary antibody directly conjugated to biotin (CR3022-biotin) for 1 hr at RT. Next, the cells were washed 3x in PBS and avidin-HRP was added for 1 hr at RT followed by three washes in PBS. Foci were visualized using TrueBlue HRP substrate (KPL, # 5510-0050) and imaged on an ELISPOT reader (CTL). Antibody neutralization was quantified by counting the number of foci for each sample using the Viridot program[2]. The neutralization titers were calculated as follows: 1 - (ratio of the mean number of foci in the presence of sera and foci at the highest dilution of respective sera sample). Each specimen was tested in duplicate. The FRNT-50 titers were interpolated using a 4-parameter nonlinear regression in GraphPad Prism v9.0.2.4.3. For samples that do not neutralize at the limit of detection at 50%, a value of 5 was plotted and used for geometric mean calculations.

### SUPPLEMENTARY METHODS

#### Propagation and Characterization of Viral Stocks

VeroE6 cells were obtained from ATCC (clone E6, ATCC, #CRL-1586). SARS-CoV-2/human/USA/GA-EHC-083E/2020) (referred to as the D614G variant) was derived from a residual nasopharyngeal swab collected from an Emory Healthcare patient in March 2020. The isolation and sequencing was previously described[3]. The B.1.351 variant isolate, kindly provided by Dr. Andy Pekosz [John Hopkins University (JHU), Baltimore, MD], was propagated once in Vero-TMPRSS2 cells to generate P2 viral stocks. Viral titers were determined by plaque assay on Vero-TMPRSS2 cells. Viral stocks were sequenced as described below and stored at -80C until use.

#### Deep Sequencing of Virus Isolate

Illumina-ready libraries were generated using NEBNext Ultra II RNA Prep reagents (New England BioLabs) as previously described[4]. Briefly, we fragmented RNA, followed by double-stranded cDNA synthesis, end repair, and adapter ligation. The ligated DNA was then barcoded and amplified by a limited cycle PCR and the barcoded Illumina libraries were sequenced by using paired-end 150-base protocol on a NextSeq 2000 (Illumina). Demultiplexed sequence reads were analyzed in the CLC Genomics Workbench v.21.0.3 by (i) trimming for quality, length, and adaptor sequence, (ii) mapping to the Wuhan-Hu-1 SARS-CoV-2 reference (GenBank accession number: NC_045512), (iii) improving the mapping by local realignment in areas containing insertions and deletions (indels), and (iv) generating both a sample consensus sequence and a list of variants. Default settings were used for all tools.

#### Titration of B.1.351 in Golden Syrian Hamsters

Golden Syrian hamsters, aged 8-9 weeks old, were randomized into groups of 10 based on weight, with each group containing a 1:1 male:female ratio. Hamsters were inoculated intranasally with total doses of $1x10^3$ – $1x10^5$ PFU of SARS-CoV-2 JHU B.1.351 P2 in a final volume of 100 µL split between each nostril. Body weight observations were made daily post-infection.

**Titration of B.1.351 in Rhesus Macaques**

Three year-old male and female Indian-origin rhesus macaques were sorted by age and weight and then stratified into groups. NHP were challenged with total doses of $2.4x10^5$ and $2.4x10^6$ PFU of JHU SARS-CoV-2 B.1.351 P2; the viral inoculum was administered in 3 mL IT and 1 mL IN (0.5 mL into each nostril). On days 2, 4, and 6 post-challenge, BAL and nasal swabs were collected and assessed for sgRNA as detailed in the below methods.

**Histopathology and Immunohistochemistry (IHC)**

As previously described [5], NHP lung tissue sections were stained with hematoxylin and eosin (H&E) for routine histopathology and a rabbit polyclonal SARS-CoV-2 (GeneTex, GTX135357) for detection of SARS-CoV-2 virus antigen. All samples were evaluated by a boarded-certified veterinary pathologist.

**Intracellular Cytokine Staining**

Cryopreserved peripheral-blood mononuclear cells were thawed, rested overnight, and stimulated with SARS-CoV-2 S protein (S1 and S2, homologous to the vaccine insert) and costimulatory antibodies anti-CD28 and anti-CD49d (clones CD28.2 and 9F10, BD Biosciences). Negative controls received an equal concentration of dimethyl sulfoxide (without peptides) and costimulatory antibodies. Cytokine staining was performed as described previously [5].

**TCID50 Quantification of SARS-CoV-2 from BAL**

Vero-TMPRSS2 cells were plated at 25,000 cells/well in Dulbecco's Modified Eagle Medium (DMEM) + 10% FBS + Gentamicin and the cultures were incubated at 37°C, 5.0% $CO_2$. Cells should be 80-100% confluent the following day. Medium was aspirated and replaced with 180 µL of DMEM + 2% FBS + gentamicin. Twenty (20) µL of BAL sample was added to top row in quadruplicate and mixed using a P200 pipettor 5 times. Using the pipettor, 20 µL was transferred to the next row, and repeated down the plate (columns A-H) representing 10-fold dilutions. The tips were disposed for each row and repeated until the last row. Positive (virus stock of known infectious titer in the assay) and negative (medium only) control wells were included in each assay setup. The plates were incubated at 37ºC, 5.0% $CO_2$ for 4 days. The cell monolayers are visually inspected for cytopathic effect (CPE). TCID50 value wass calculated using the Read-Muench formula. For optimal assay performance, the TCID50 value of the positive control should test within 2-fold of the expected value.

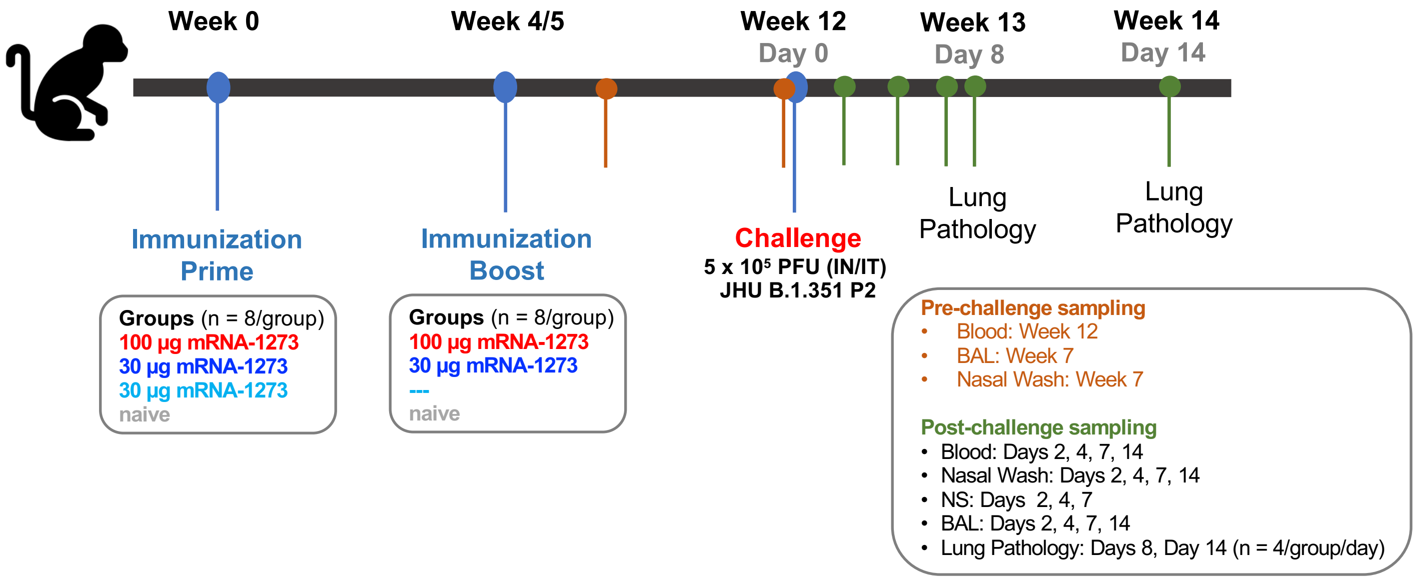

**Figure S1. Study design: Ability of mRNA-1273 to protect NHP against B.1.351 challenge.** Nonhuman primates (NHP), Rhesus macaques (n = 8/group), were immunized with mRNA-1273 on the following schedule: Group 1: 0 and 4 weeks, 100 µg; Group 2: 0 and 5 weeks, 30 µg; Group 3: week 0, 30 µg. Naïve aged-matched NHP were included as controls. At week 12, animals were challenged with a total of $5x10^5$ PFU of SARS-CoV-2 B.1.351. The viral inoculum was administered as $3.75x10^5$ PFU in 3 mL intratracheally (IT) and $1.25x10^5$ PFU in 1 mL intranasally (IN) in a volume of 0.5 mL into each nostril. Sera were collected at weeks 7 and week 12. Bronchoalveolar lavages (BAL) and nasal washes were also collected at week 7. Sera, BAL, and nasal washes were collected post-challenge on days 2, 4, 7, and 14, as indicated. Lung pathology was assessed on day 8 post-challenge in a subset of animals (n = 4/group).

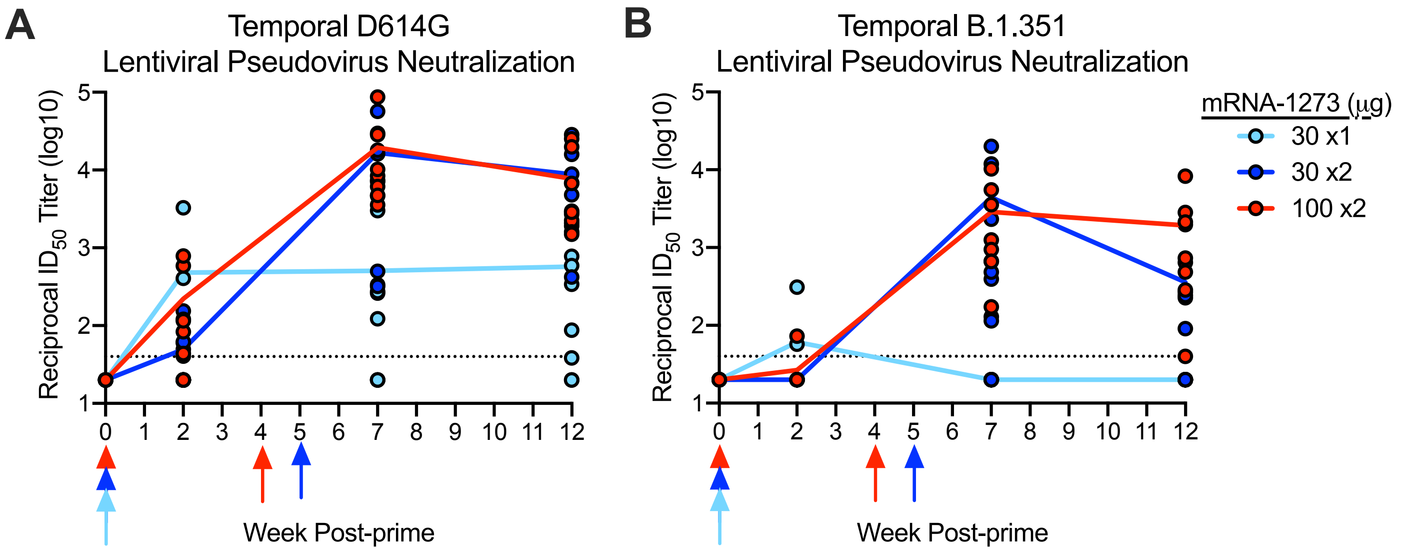

**<u>Figure S2</u>. Temporal neutralizing antibody responses following mRNA-1273 immunization.** Rhesus macaques were immunized according to Figure S1. Sera collected at weeks 0, 2, 7, and 12 were assessed for SARS-CoV-2 D614G (A), and B.1.351 (B) lentiviral-based pseudovirus neutralization. Circles represent individual NHP and may overlap where values are equal; lines represent geometric mean titers (GMT). Dotted lines indicate neutralization assay limits of detection. Arrows point to immunization weeks.

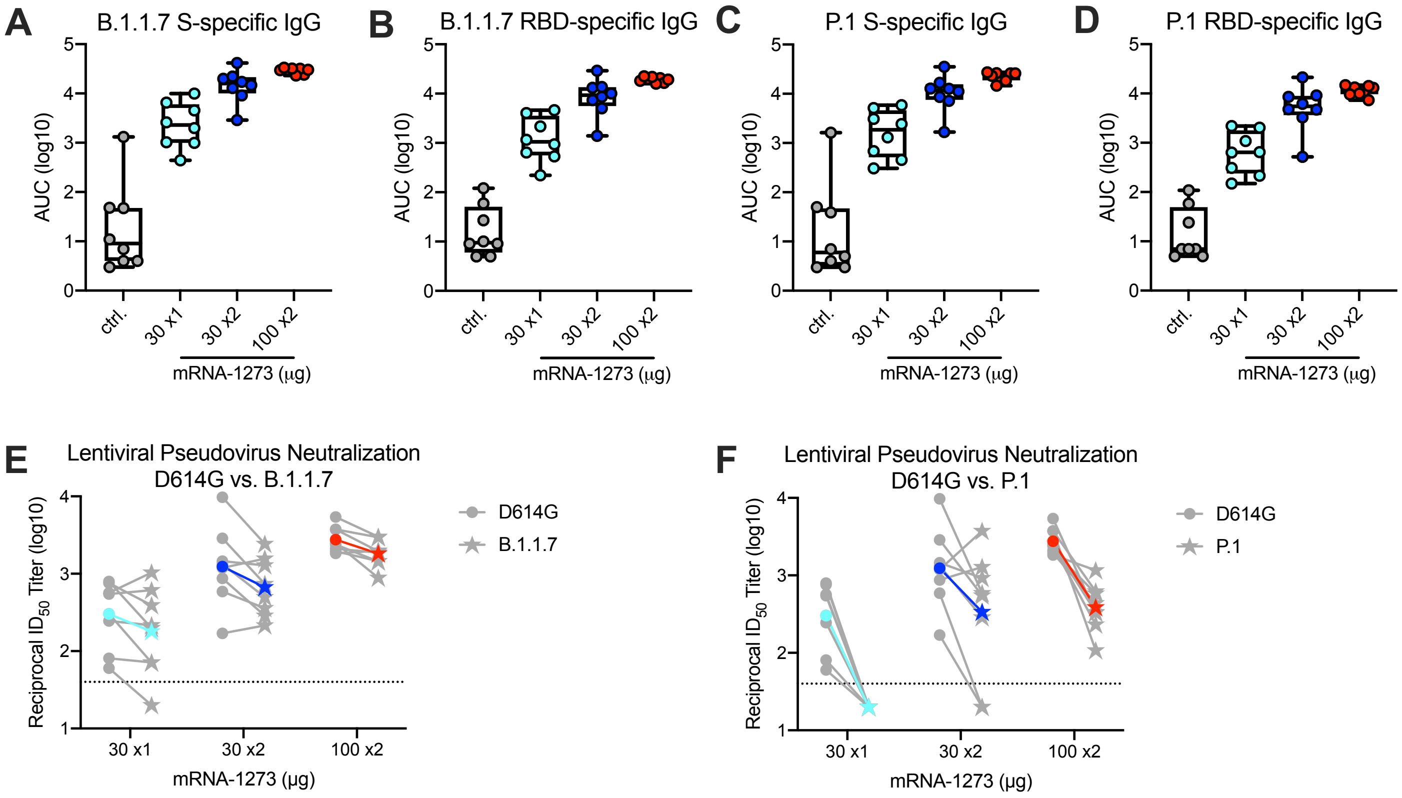

**<u>Figure S3</u>. Cross-reactivity of mRNA-1273-immune NHP serum antibodies to global variants.** Rhesus macaques were immunized according to Figure S1. Sera collected at week 12 were assessed for SARS-CoV-2 B.1.1.7 (A-B) and P.1 (B-C) S-specific (A, C) and RBD-specific (B, D) IgG by MULTI-ARRAY ELISA, and SARS-CoV-2 D614G (E-F), B.1.1.7 (E), and P.1 (F) lentiviral-based pseudovirus neutralization. (A-D) Circles represent individual NHP. Boxes and horizontal bars denote the IQR and medians, respectively; whisker end points are equal to the maximum and minimum values. (E-F) Gray lines represent individual NHP, and colored lines represent GMT. Dotted lines indicate neutralization assay limits of detection.

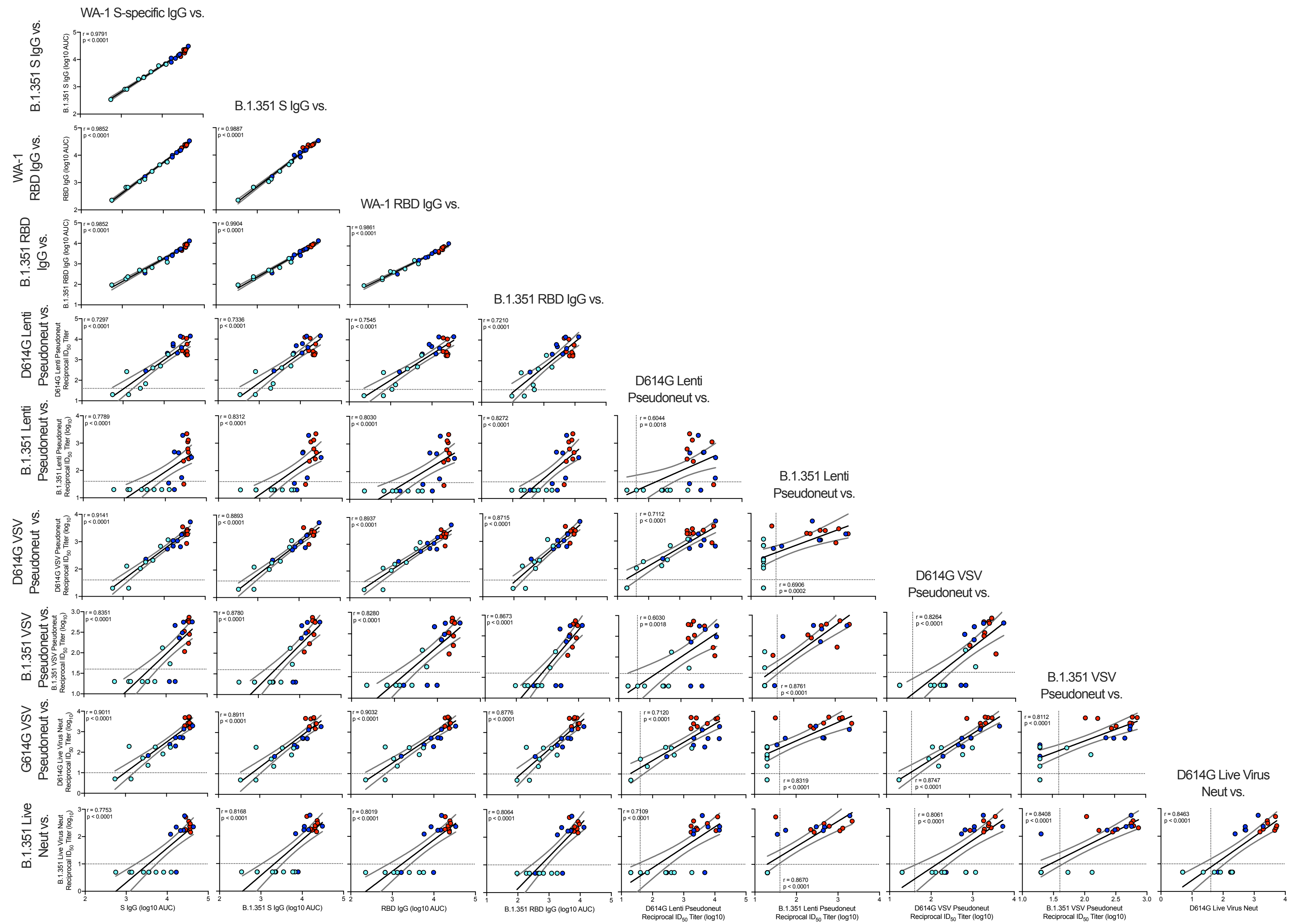
WA-1 S-specific IgG vs.
B.1.351 S IgG vs.
WA-1 RBD IgG vs.
B.1.351 RBD IgG vs.
D614G Lenti Pseudoneut vs.
B.1.351 Lenti Pseudoneut vs.
D614G VSV Pseudoneut vs.
B.1.351 VSV Pseudoneut vs.
D614G Live Virus Neut vs.
B.1.351 S IgG vs.
WA-1 RBD IgG vs.
B.1.351 RBD IgG vs.
D614G Lenti Pseudoneut vs.
B.1.351 Lenti Pseudoneut vs.
D614G VSV Pseudoneut vs.
B.1.351 VSV Pseudoneut vs.
G614G VSV Pseudoneut vs.
B.1.351 Live Neut vs.
S IgG (log10 AUC)
B.1.351 S IgG (log10 AUC)
RBD IgG (log10 AUC)
B.1.351 RBD IgG (log10 AUC)
D614G Lenti Pseudoneut Reciprocal $ID_{50}$ Titer (log10)
B.1.351 Lenti Pseudoneut Reciprocal $ID_{50}$ Titer (log10)
D614G VSV Pseudoneut Reciprocal $ID_{50}$ Titer (log10)
B.1.351 VSV Pseudoneut Reciprocal $ID_{50}$ Titer (log10)
D614G Live Virus Neut

**Figure S4**. **Correlations of humoral antibody analyses.** Rhesus macaques were immunized according to Figure S1C. Plots show correlations between SARS-CoV-2 WA-1 S-specific IgG, B.1.351 S-specific IgG, WA-1 RBD-specific IgG, B.1.351 RBD-specific IgG, D614G lentiviral-based pseudovirus neutralization, B.1.351 lentiviral-based pseudovirus neutralization, D614G VSV-based pseudovirus neutralization, B.1.351 VSV-based pseudovirus neutralization, D614G focus reduction neutralization, and B.1.351 focus reduction neutralization at week 12. Circles represent individual NHP, where colors indicate mRNA-1273 dose as defined in Figure S1. Dotted lines indicate assay limits of detection. Black and gray lines indicate linear regression and 95% confidence interval, respectively. 'r' and 'p' represent Spearman's correlation coefficients and corresponding p-values, respectively.

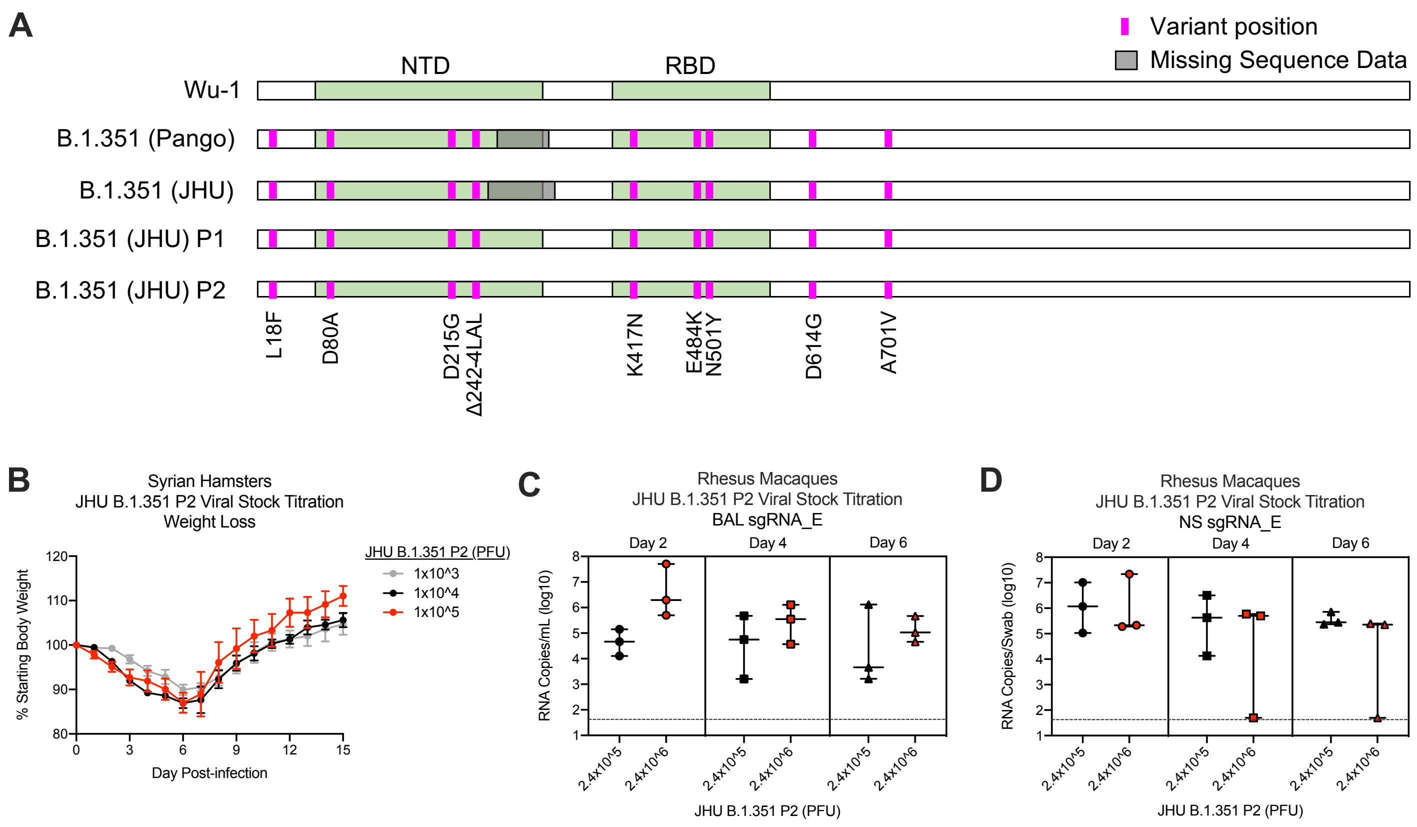

**<u>Figure S5</u>. Characterization of B.1.351 viral isolate.** A SARS-CoV-2 B.1.351 clinical isolate was first passaged (P1) at Johns Hopkins University (JHU) on Vero cells then passaged again (P2) on Vero/TMPRSS2 cells. P1 and P2 underwent shotgun deep sequencing. (A) Alignment of S protein consensus sequence, where pink and gray indicate variant amino acid position and missing sequence data, respectively. (B) Syrian hamsters (n = 10/group) were infected with $1x10^3$ (gray), $1x10^4$ (black), or $1x10^5$ (red) dilution of JHU B.1.351 P2 and monitored for weight loss for 15 days post-infection. Circles and error bars represent means and SEM, respectively. (C-D) Rhesus macaques (n = 3/group) were infected with $2.4x10^5$ (black) or $2.4x10^6$ (red) PFU of JHU B.1.351 P2 and viral replication was assessed by detection of SARS-CoV-2 E-specific sgRNA in BAL (C) and NS (D) on days 2, 4, and 6 post-infection. Boxes and horizontal bars denote the interquartile ranges (IQR) and medians, respectively; whisker end points are equal to the maximum and minimum values. Dotted lines indicate assay limits of detection.

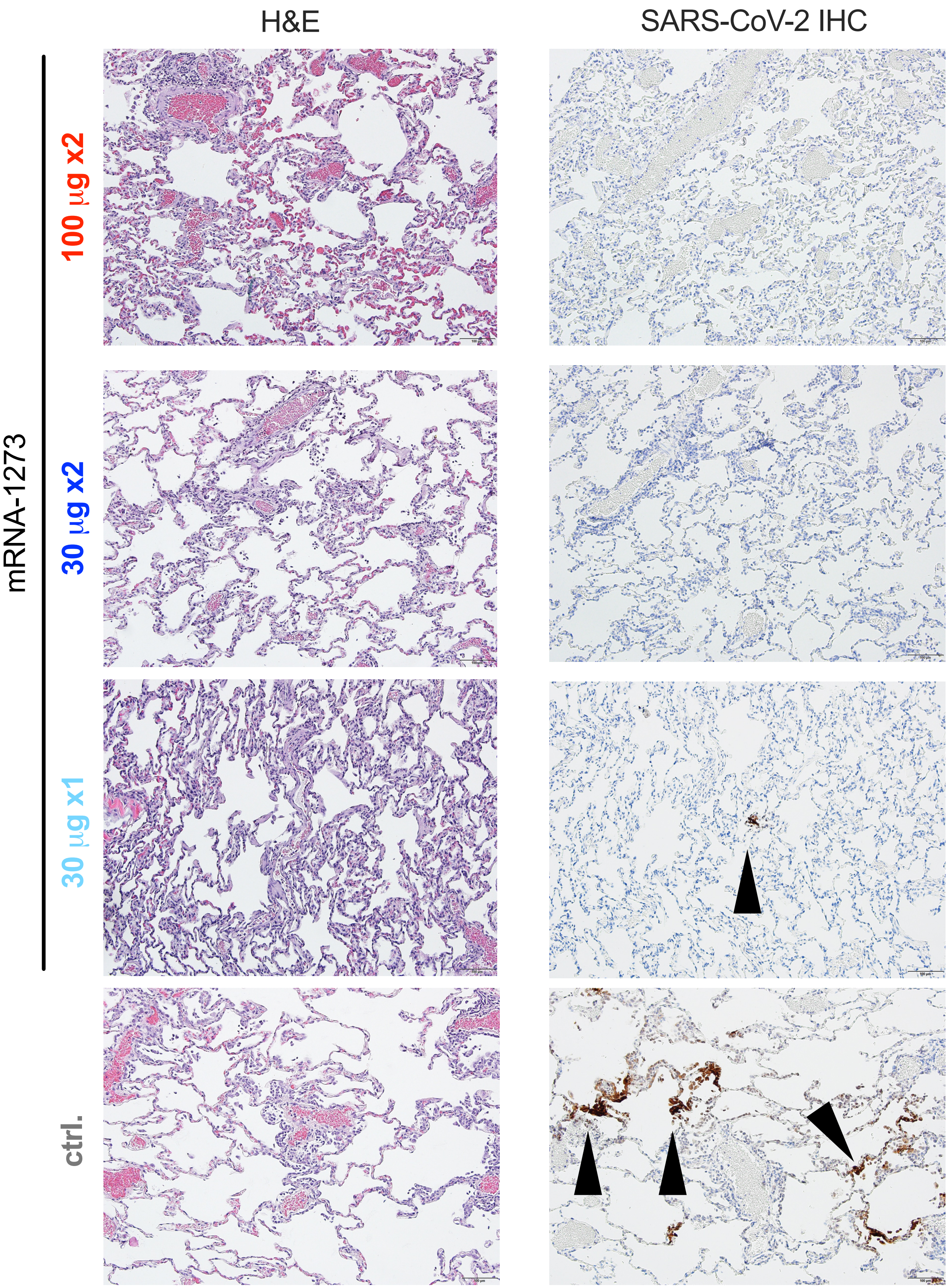
H&E
SARS-CoV-2 IHC
mRNA-1273
100 µg x2
30 µg x2
30 µg x1
ctrl.

**Figure S6. Post-challenge lung histopathological analysis and viral detection.** Rhesus macaques were immunized and challenged as described in Figure S1. Eight days post-challenge, lung samples (n = 4/group) were evaluated for the presence of inflammation by hematoxylin and eosin (H & E) staining (left) and evidence of virus infection by immunohistochemistry (IHC) for SARS-CoV-2 viral antigen (right). Representative images show the location and distribution of SARS-CoV-2 viral antigen in serial lung tissue sections. Arrows indicate areas positive for viral antigen. Each image is taken at 10x magnification; scale bars represent 100 microns.

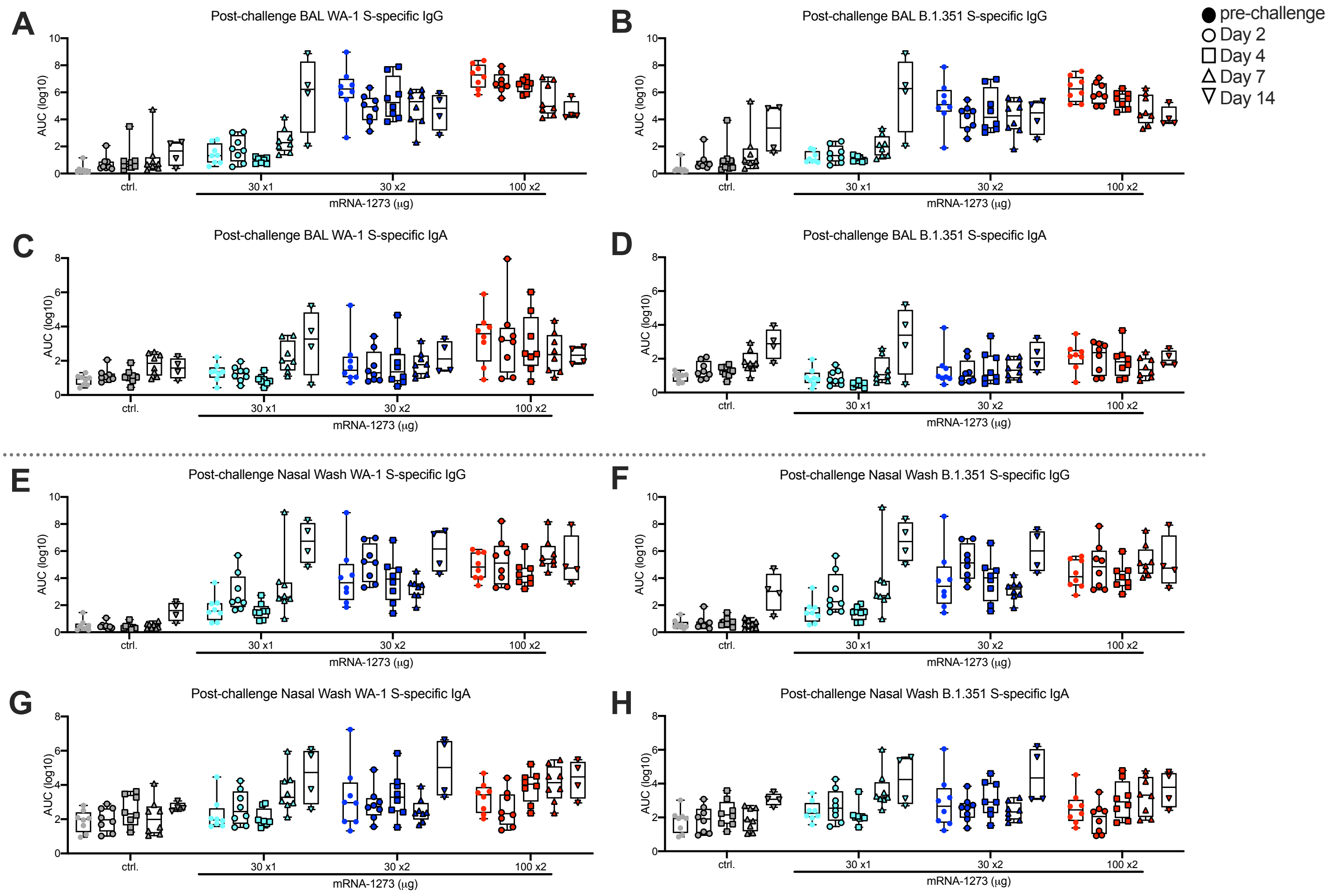

**<u>Figure S7</u>. Mucosal antibody responses following SARS-CoV-2 challenge in mRNA-1273-immunized NHP.** Rhesus macaques were immunized and challenged according to Figure S1. BAL (A-D) and nasal washes (E-H) collected at week 7 (filled circles) and days 2 (circles), 4 (squares), 7 (triangles), and 14 (inverted triangles) post-challenge were assessed for SARS-CoV-2 WA-1 (A, C, E, G) and B.1.351 (B, D, F, H) S-specific IgG (A-B, E-F) and IgA (C-D, G-H) by MULTI-ARRAY ELISA. Boxes and horizontal bars denote the IQR and medians, respectively; whisker end points are equal to the maximum and minimum values. Symbols represent individual NHP.

| mRNA-1273 (µg) | Animal ID | Inflammation (H&E)[1] | | | IHC[5] | | |
|---|---|---|---|---|---|---|---|
| | | Lc[2] | Rmid[3] | Rc[4] | Lc | Rmid | Rc |
| **100 x2** | 08N012 | - | +/- | - | - | - | - |
| | DG3T | + | +/- | +/- | - | - | - |
| | 15C228 | - | - | - | - | - | - |
| | DGEH | + | +/- | +/- | - | - | - |
| **30 x2** | 35987 | +/- | +/- | + | - | - | - |
| | DGB3 | - | +/- | ++ | - | - | - |
| | MB13 | - | +/- | +/- | - | - | - |
| | DH49 | - | - | +/- | - | - | - |
| **30 x1** | 35975 | - | +/- | +/- | - | +/- | - |
| | 17C179 | - | - | +/- | - | - | - |
| | DGI2 | +/- | +/- | +/- | - | +/- | - |
| | DGJH | - | +/- | +++ | - | - | - |
| **Naïve** | G25M | + | + | ++ | + | +/- | +/- |
| | 15C024 | + | ++ | ++ | - | + | + |
| | DFZi | + | ++ | ++ | - | + | + |
| | A6V055 | ++ | ++ | ++ | ++ | + | + |

**Table S1. Assessment of lung inflammation and viral antigen on day 8 following SARS-CoV-2 B.1.351 challenge in mRNA-1273-immunized NHP.** Lung tissue was evaluated for the presence of inflammation by (H & E staining and evidence of virus infection by IHC for SARS-CoV-2 viral antigen.

[1] Inflammation scoring: - = minimal to absent, +/- = minimal to mild, + = mild to moderate, ++ = moderate to severe, +++ = severe
[2] Lc: left caudal lung lobe
[3] Rmid: right middle lung lobe
[4] Rc: right caudal lung lobe
[5] Immunohistochemistry SARS-CoV-2 antigen (Ag): - = no detection of virus Ag, +/- = rare/occasional Ag+ foci, + = multiple Ag+ foci

| Group | Animal ID | WA-1 S-specific IgG[1] (IU/mL)[2] | WA-1 RBD-specific IgG (IU/mL) |
|---|---|---|---|
| **100 µg mRNA-1273 x2** | 08N012 | 1002.4 | 2069.5 |
| | 1F-8 | 734.0 | 1255.8 |
| | DG3T | 1161.7 | 2329.0 |
| | 15C228 | 1254.9 | 1969.4 |
| | DG94 | 1203.4 | 2160.1 |
| | DGMZ | 719.7 | 1268.8 |
| | DGEH | 1285.4 | 3181.5 |
| | DGBAA | 1228.0 | 2206.4 |
| | **GMT** | **1049.0** | **1972.7** |
| **30 µg mRNA-1273 x2** | 35987 | 76.6 | 143.6 |
| | HNM | 781.5 | 1318.5 |
| | DGB3 | 491.6 | 879.4 |
| | MI03 | 2392.3 | 4460.5 |
| | DH26 | 777.4 | 1454.7 |
| | MB13 | 434.5 | 621.4 |
| | DH49 | 542.6 | 1062.8 |
| | 36845 | 279.7 | 460.8 |
| | **GMT** | **495.1** | **870.2** |
| **30 µg mRNA-1273 x1** | 35975 | 75.4 | 110.2 |
| | HZF | 284.7 | 451.9 |
| | LV41 | 97.8 | 170.9 |
| | 17C179 | 10.1 | 18.7 |
| | DGF1 | 21.1 | 45.5 |
| | DGW2 | 195.3 | 283.2 |
| | DGI2 | 28.3 | 42.3 |
| | DGJH | 54.1 | 74.0 |
| | **GMT** | **58.3** | **94.6** |
| **Naïve** | G25M | 1.2 | 5.9 |
| | 07N001 | *0.3*[3] | *1.6*[4] |
| | 15C184 | 1.4 | 10.1 |
| | 15C024 | *0.3* | *1.6* |
| | DGEI | *0.3* | *1.6* |
| | A12V163 | *0.3* | *1.6* |
| | DFZi | *0.3* | *1.6* |
| | A6V055 | 36.1 | *1.6* |
| | **GMT** | **0.8** | **2.4** |

**Table S2. SARS-CoV-2 WA-1 S- and RBD-specific antibody responses in international units (IU).**

[1]Sera were collected at week 12 and assessed for SARS-CoV-2 WA-1 S- and RBD-specific IgG by 4-plex MULTI-ARRAY ELISA.
[2]IgG quantities were extrapolated as IU/mL using WHO standards.
[3]S-specific IgG lower limit of detection = 0.3076 IU/mL
[4]RBD-specific IgG lower limit of detection = 1.5936 IU/mL.
